# High throughput functional profiling of genes at intraocular pressure loci reveals distinct networks for glaucoma

**DOI:** 10.1101/2023.07.10.548340

**Authors:** Connor J Greatbatch, Qinyi Lu, Sandy Hung, Kristof Wing, Helena Liang, Xikun Han, Tiger Zhou, Owen M Siggs, David A Mackey, Anthony L Cook, Anne Senabouth, Guei-Sheung Liu, Jamie E Craig, Stuart MacGregor, Joseph E Powell, Alex W Hewitt

**Author notes:** These authors contributed equally to this work (Connor J Greatbatch, Qinyi Lu). These authors jointly supervised this work (Jamie E Craig, Stuart MacGregor, Joseph E Powell, Alex W Hewitt). Address for Correspondence: Professor Alex Hewitt, Menzies Institute for Medical Research University of Tasmania, 17 Liverpool St, Hobart, TAS, 7000.

## Abstract

**INTRODUCTION:** Primary open angle glaucoma (POAG) is a leading cause of blindness globally. Characterised by progressive retinal ganglion cell degeneration, the precise pathogenesis remains unknown. Genome-wide association studies (GWAS) have uncovered many genetic variants associated with elevated intraocular pressure (IOP), one of the key risk factors for POAG. This study sought to investigate the morphological and transcriptional consequences of perturbation of key genes at IOP loci in trabecular meshwork cell (TMC); the cellular regulators of IOP. We aimed to identify genetic and morphological variation that can be attributed to TMC dysfunction and raised IOP in POAG.

**METHODS:** 62 genes across 55 loci were knocked-out in a primary human TMC line. Each knockout group, including five non-targeting control groups, underwent single-cell RNA-sequencing (scRNA-seq) for differentially-expressed gene (DEG) analysis. Multiplexed fluorescent staining of key organelles, was coupled with high-throughput microscopy for single-cell morphological analysis using CellProfiler image analysis.

**RESULTS:** Across many of the individual gene knockouts scRNA-seq highlighted genes relating to matrix metalloproteinases and interferon-induced proteins. Our work has prioritised genes at four loci of interest to identify gene knockouts that may contribute to the pathogenesis of POAG, including *ANGPTL2, LMX1B, CAV1,* and *KREMEN1*. Three genetic networks of gene knockouts with similar transcriptomic profiles were identified (*ABO* / *CAV1* / *MYOC*, *ANGPT2* / *PKHD1* / *TNS1* / *TXNRD2*, and *CAPZA1* / *KALRN* / *LMO7* / *PLEKHA7* / *GNB1L* / *TEX41*), suggesting a synergistic function in trabecular meshwork cell physiology. *TEK* knockout caused significant upregulation of nuclear granularity on morphological analysis, whilst knockout of *TRIOBP, TMCO1* and *PLEKHA7* increased granularity and intensity of actin and the cell-membrane.

**CONCLUSION:** High throughput analysis of cellular structure and function through multiplex fluorescent single-cell analysis and scRNA-seq assays enabled the direct study of genetic perturbations at the single-cell resolution. This work provides a framework for investigating the role of genes in the pathogenesis of glaucoma and heterogenous diseases with a strong genetic basis.

## INTRODUCTION

Glaucoma is a heterogeneous group of diseases leading to irreversible blindness with characteristic optic nerve damage. The most common glaucoma subtype is primary open-angle glaucoma (POAG).^1, 2^ Elevated intraocular pressure (IOP) is the only known modifiable risk factor and plays a major role in the progression of POAG. The circulatory system maintains IOP in the anterior segment of the eye ^3, 4^ Aqueous humor is produced by the ciliary body and passes through the pupil before draining out to the episcleral blood vessels via conventional or unconventional pathways.^1, 5^ The conventional outflow pathway through the trabecular meshwork accounts for approximately 80% of total aqueous humor outflow. Structural alterations observed in the trabecular meshwork are considered to increase outflow resistance in POAG.^4, 6, 7^

Many POAG-associated loci have been identified through genome-wide association studies (GWAS), with loci encompassing Caveolin 1 and 2 (*CAV1/CAV2*), Transmembrane and coiled-coil domain-containing protein 1 (*TMCO1*), cyclin-dependent kinase inhibitor 2B antisense RNA 1 (*CDKN2B-AS1*), ATB binding cassette subfamily A member 1 (*ABCA1*), actin filament associated protein 1 (*AFAP1*), GDP-mannose 4,6-dehydratase (*GMDS*), Forkhead Box C1 (*FOXC1*), thioredoxin reductase 2 (*TXNRD2*), and Ataxin 2 (*ATXN2*).^9–12^ Furthermore, protein altering variants in genes such as *MYOC, LTBP2, FOXC1, GMDS* and *CYP1B1* have been found to cause both congenital and juvenile onset glaucoma. These particular variants are generally associated with abnormal development of the aqueous circulatory system and Schlemm’s canal rather than maintenance, however some are also involved in maintenance such as *TEK.*^13–17^

More recently, a GWAS meta-analysis identified 85 novel SNPs associated with IOP using data from the UK Biobank, the International Glaucoma Genetic Consortium, and the Australian & New Zealand Registry of Advanced Glaucoma Cohort.^18^ Novel gene variants, including *ANGPT1, ANKH, MECOM* and *ETS1* were associated with POAG and IOP. However, this study also identified SNPs at *ADAMTS6*, *MYOF, ANAPC1, GLIS3*, and *FNDC3B* that are associated with phenotypes such as central corneal thickness and corneal hysteresis.^18^ This highlights potential confounding factors in GWAS that make identification of genes implicated in the pathogenesis of POAG challenging. Furthermore, various SNPs identified in IOP associated GWAS are associated with more than one gene, making it difficult to precisely implicate the disease-causative gene.

Recent advances in clustered regularly interspaced short palindromic repeats (CRISPR) and single-cell RNA sequencing (scRNA-seq) technology have allowed for high-throughput genetic screens at single-cell transcriptome resolution. In CRISPR droplet sequencing (CROP-seq), a guide-RNA (gRNA)-encoding vector makes gRNAs detectable in scRNA-seq, and as such, these gRNAs can be used to tag individual cells.^19^ To investigate the role of POAG-associated loci in TMCs, we knocked out gene candidates in human TMC lines using CROP-seq. We then performed single-cell RNA sequencing as well as morphological profiling to identify the genotypic and phenotypic roles of each gene. The cell painting protocol involves cultured cells being stained with fluorescent dyes to reveal eight cellular substructures, thus allowing morphological features to be extracted from individual cells to display the effects of genetic perturbation.^20, 21^ Morphological profiling can then be undertaken using CellProfiler, a high-throughput single-cell image analysis program designed to extract and analyze over one thousand phenotypic features. Taken together, this study screens gene candidates based on expression profiles and morphology profiles and helps understand the pathway in which these genes are involved in the causation of elevated IOP in TMCs. **(Figure 1).**

**Figure 1:**
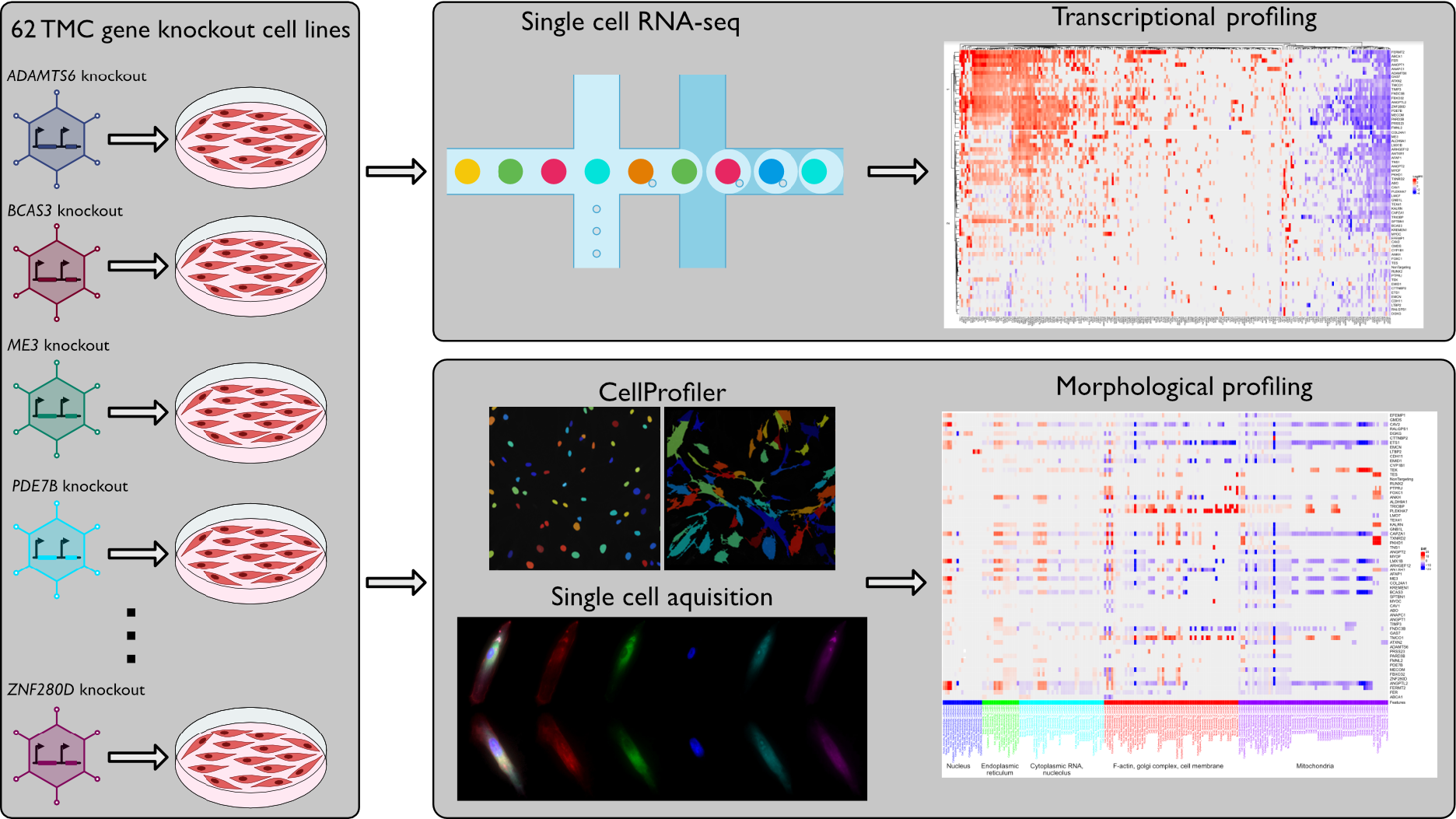
Schematic Overview of the Study. Primary human TMC lines were cultured and split into 62 groups, each with a single gene knockout at IOP-associated loci. A further 5 cell lines were maintained as control groups. All groups underwent transcriptional profiling and morphological profiling analysis to identify the transcriptomic and morphological effects of particular gene knockouts. This demonstrated genes that may play a role in the pathogenesis of POAG, identified novel genetic networks in IOP regulation as well as prioritising genes at multi-gene loci.

## RESULTS

### Data overview

A total of 105,273 cells were captured with 25,879 (24.58%) cells passing quality control filtering to be included in transcriptional profiling. Differentially-expressed gene (DEG) analysis was performed to investigate the effects of gene knockout at select IOP-associated loci. The Euclidean distance of DEG expression between each knockout group and controls was computed to identify gene knockouts with similar expression patterns that may indicate novel genetic networks involved in the pathogenesis of POAG. DEG analysis was also used to prioritise multi-gene loci to identify a pathological variant. Ward’s hierarchical clustering method was then used to generate a cluster tree for further analysis of genetic networks (**Figure 2**). Gene knockout clusters were allocated based on branch thirty-two of the cluster tree whereby the congenital (developmental) glaucoma genes were grouped with normal controls. This is because many of the congenital glaucoma genes are hypothesised to primarily affect the development of trabecular meshwork tissue rather than its maintenance which is affected in POAG.^22–24^.

**Figure 2:**
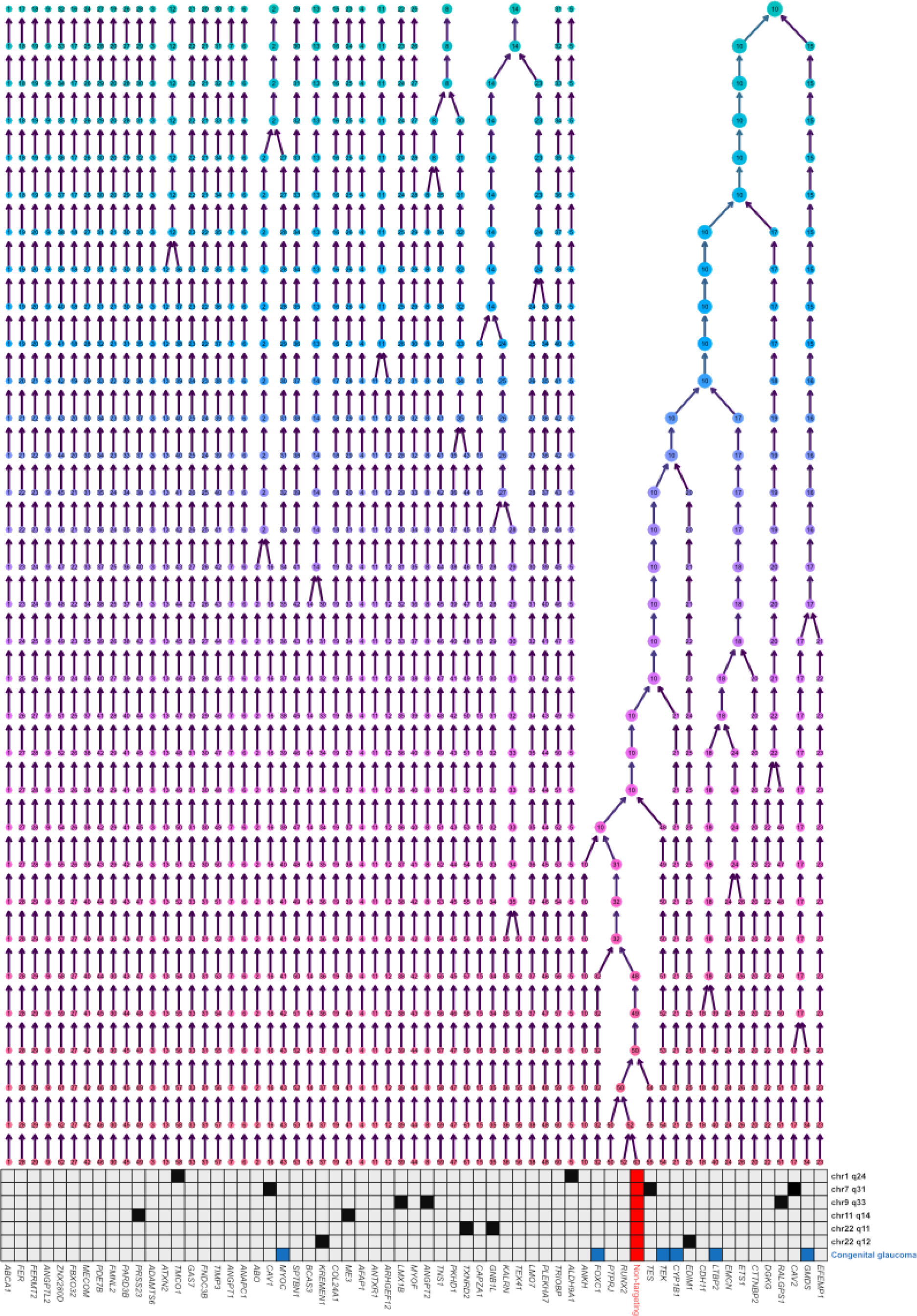
Wards cluster tree displaying the hierarchical clustering of each cell line based on the RNA expression profiles. The method of ward.D2 was applied, the distance between each group within one branch was closer than those located in different branches. The bottom grid shows the multi-gene loci groups, the potential positive genes (blue) associated with congenital glaucoma, and the non-targeting control group (red).

### Key up- and downregulated DEGs of interest

DEG analysis revealed key genetic families that may play a role in the pathogenesis of POAG. Volcano plots of all gene knockouts are displayed in **Supplementary Figure 1**. Matrix metalloproteinases were widely upregulated across many of the gene knockouts. *MMP1* was upregulated in 71% (44/62) of knockout groups and downregulated in 9% (6/62). *MMP3* was similarly upregulated in 61% (38/62) and downregulated in 5% (3/62). Finally, *MMP10* was upregulated in 30% (19/62) of the knockout groups. The proteins encoded by these genes are part of a family of proteins involved in the breakdown of extracellular matrix in physiologic and pathologic processes. The matrix metalloproteinase family of proteins have also been previously implemented in TMC function and the pathogenesis of POAG, with upregulation of MMPs 1, 9, and 12 associated with POAG.^25–27^ Another group of highly upregulated DEGs were interferon-alpha (*IFI27, IFI6)* and interferon-induced proteins (*IFI44L, IFIH1, IFIT1, IFIT2, IFIT3, IFITM1, IFIM10*). These genes were upregulated in 40 - 80% (25/62 - 50/62) of the knockout groups **(Figure 3).** These proteins are all generally involved in antiviral immunity, however, there have been minimal direct associations between glaucoma and interferon-related proteins.

**Figure 3:**
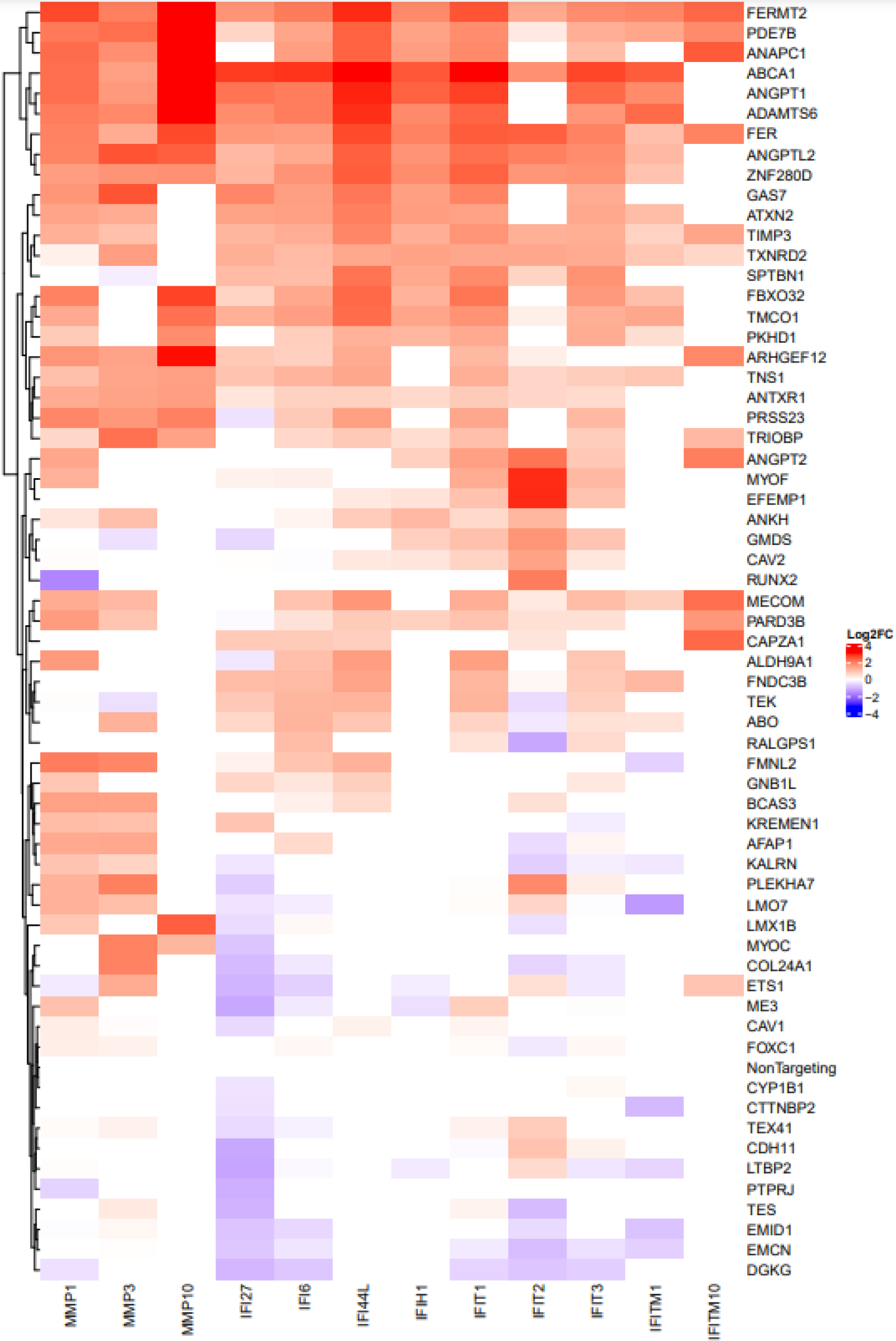
Heatmap illustrating significant up-regulation of matrix metalloproteinases (*MMP1, MMP3, MMP10*) and interferon-related proteins (*IFI27, IFI6, IFI44L, IFIH1, IFIT1, IFIT2, IFIT3, IFITM1, IFITM10*).

### scRNAseq clustering allows prioritisation at multi-gene loci

DEG and morphological profiling analysis was used to prioritise the most likely pathological gene at multi-gene loci **(Table 1)**. On chromosome nine, there are four loci of interest which involve *ANGPTL2, RALGPS1* and *LMX1B*. The *ANGPTL2* knockout group was shown to have the highest Euclidean distance (18.79) as well as cluster independently from control groups. Furthermore, *ANGPTL2* also had the highest number of significant DEGs (23) which were primarily involved in interferon alpha/beta signalling. The *LMX1B* knockout group had the next highest Euclidean distance (14.06) and significant DEGs (11) which are primarily involved in regulating cell proliferation. *LMX1B* also clustered independently from the non-targeting control groups. Finally, the *RALGPS1* knockout group showed a much lower Euclidean distance (7.1) as well as the lowest number of DEGs (4) whilst also clustering with the non-targeting controls’ gene expression profile. Furthermore, the morphological profiling data revealed that *RALGPS1* had almost no significant changes in cellular morphology and was concordantly clustered with non-targeting controls’ gene expression profile. The *ANGPTL2* knockout, however, evoked a significant reduction of intensity and granularity in both the mitochondrial and actin/cell membrane channels. As well as this, the *LMX1B* knockout also induced a significant reduction in morphological intensity and granularity across the mitochondrial and actin/cell membrane channels. **(Figure 4A)**.

**Figure 4:**
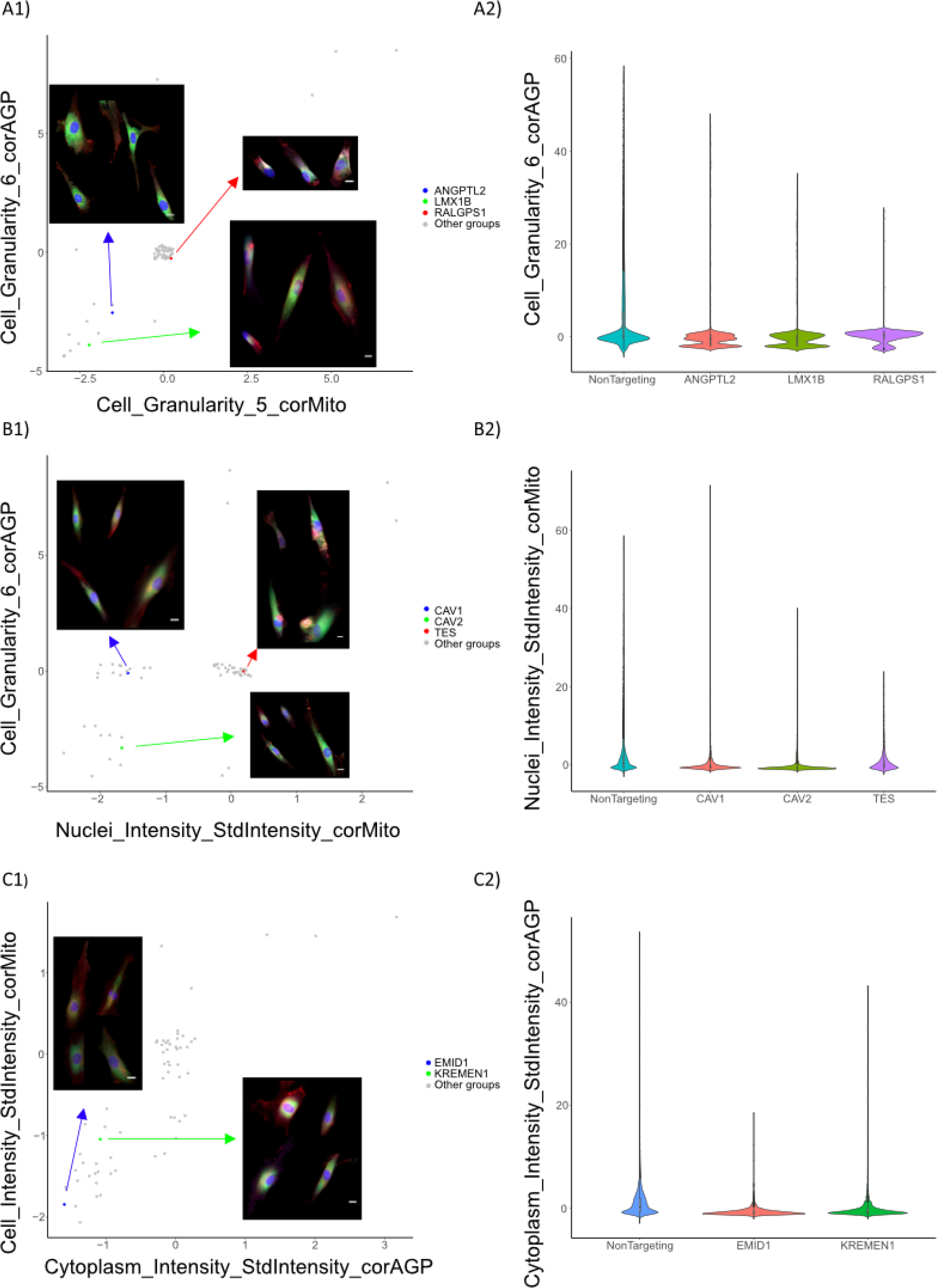
Cell features of selected genes at multi-gene loci: A: *ANGPTL2*, *LMX1B*, and *RALGPS1*; B: *CAV1*, *CAV2*, and *TES*; C: *EMID1*, and *KREMEN1*. Lane 1: Scatter plots of the mean value of each group of features (Jitter was applied to avoid overplotting). Lane 2: The distribution of value of features of each group. Scale bar: 5μm

**Table 1:**
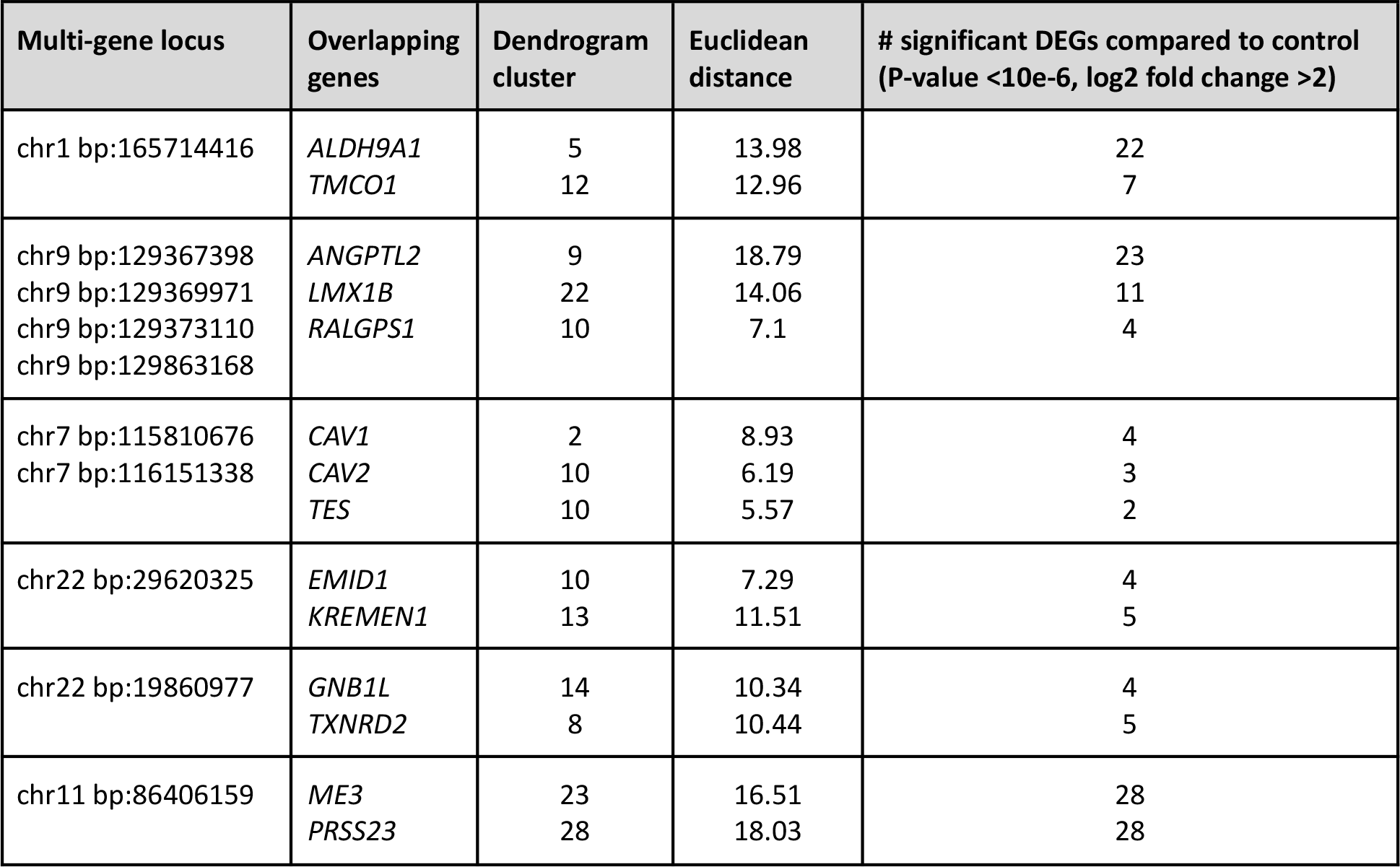
Breakdown in number of differentially expressed genes from the individual knockout of genes at overlapping loci.

On chromosome seven three genes were involved at a locus of interest; *CAV1, CAV2,* and *TES. CAV1* was the only gene to cluster independently from normal controls whilst also demonstrating a higher Euclidean distance and a slightly higher number of DEGs (4 vs 3 vs 2, respectively). The DEGs in the *CAV1* group did not cluster into a single transcriptional pathway. Curiously, the *CAV2* knockout was found to significantly upregulate the expression of *MYOC* (encoding the myocilin protein) **(Figure 5)**. Myocilin is one of the most well-evidenced pathological factors contributing to the development of early-onset POAG.^13, 28–31^ This could infer that the knockout of *CAV2* may induce cellular effects similar to *MYOC* mutations. The *CAV1* knockout resulted in a small reduction in mitochondrial intensity. The *CAV2* knockout produced a significant reduction in the intensity and granularity of the mitochondrial and actin/cell membrane channels. The *TES* knockout produced minimal morphological change as supported by clustering with the non-targeting control group **(Figure 4B)**. Finally, on chromosome 22, a multi-gene locus of interest included *EMID1* and *KREMEN1*. *KREMEN1* clustered separately to controls, unlike *EMID1,* resulting in a higher Euclidean distance and a slightly higher number of DEGs (5 vs 4). The DEGs induced by *KREMEN1* knockout also did not cluster into a particular genetic pathway. However, the *EMID1* knockout produced slightly more morphological variation, primarily as intensity reduction in the mitochondrial and actin/cell membrane channels **(Figure 4C)**. The remaining multi-gene loci all clustered independently from the control group and had similar degrees of DEG expression, which makes it difficult to resolve the prioritised gene.

**Figure 5:**
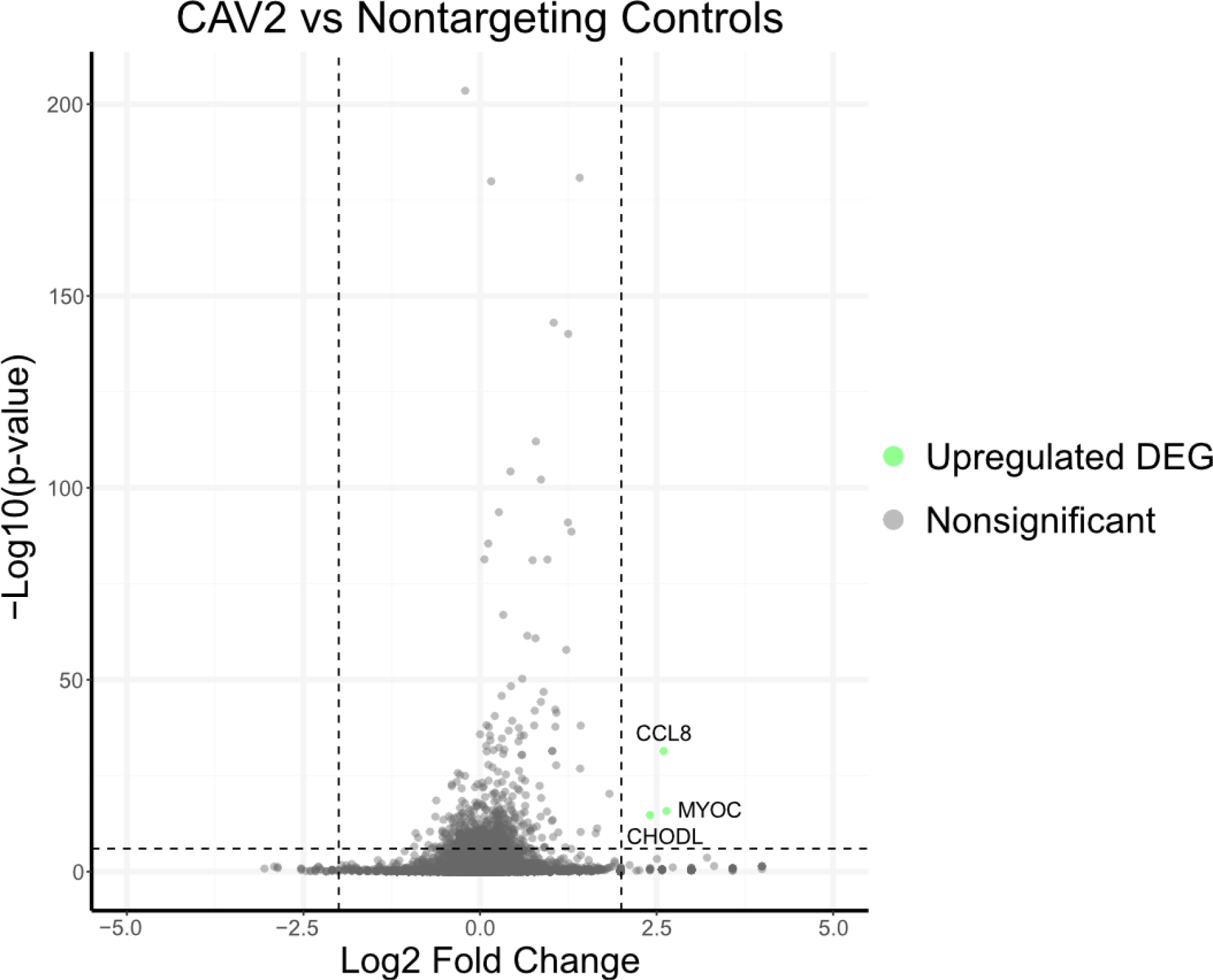
*CAV2* knockout differentially expressed genes *CAV2* knockout evoked a significant upregulation of myocilin (encoded by *MYOC*). Significant upregulation defined as Log_2_ Fold Change greater than 2 and a p-value less than 10^-6^. There were no significantly downregulated genes in the *CAV2* knockout transcriptional analysis.

### Identification of putative genetic networks involved in the pathogenesis of POAG

Gene expression of normal control cells was analysed in a genetic network which highlighted 30 (58%) of the target genes were normally expressed in the control cells. This illustrates that many of these genes play a role in TMC functioning. **(Figure 6).** Cluster analysis was performed to identify any novel genetic networks that may be involved in the pathogenesis of POAG. Using Ward’s method of hierarchical clustering, we were able to show clusters of multiple genes with similar DEG profiles. Cluster two contained three genes that had a similar DEG profile; *ABO, CAV1,* and *MYOC. MYOC* encodes myocilin and is one of the most well-known genetic causes of POAG and is highly evidenced in the literature. *CAV1* encodes for caveolin 1, which is involved in cell membrane structure and has also been speculated to regulate adhesion, endocytosis, and autophagy in TM cells.^32^ *ABO* encodes for proteins that determine blood group has been speculated to be involved in POAG however the exact mechanisms remain unknown. Each of these gene knockouts invoked several DEGs, however, the most common were the upregulation of *TAC1* and *LCE1C*. *TAC1* (Tachykinin precursor 1) encodes for peptides involved in neuronal excision and potent vasodilation.

**Figure 6:**
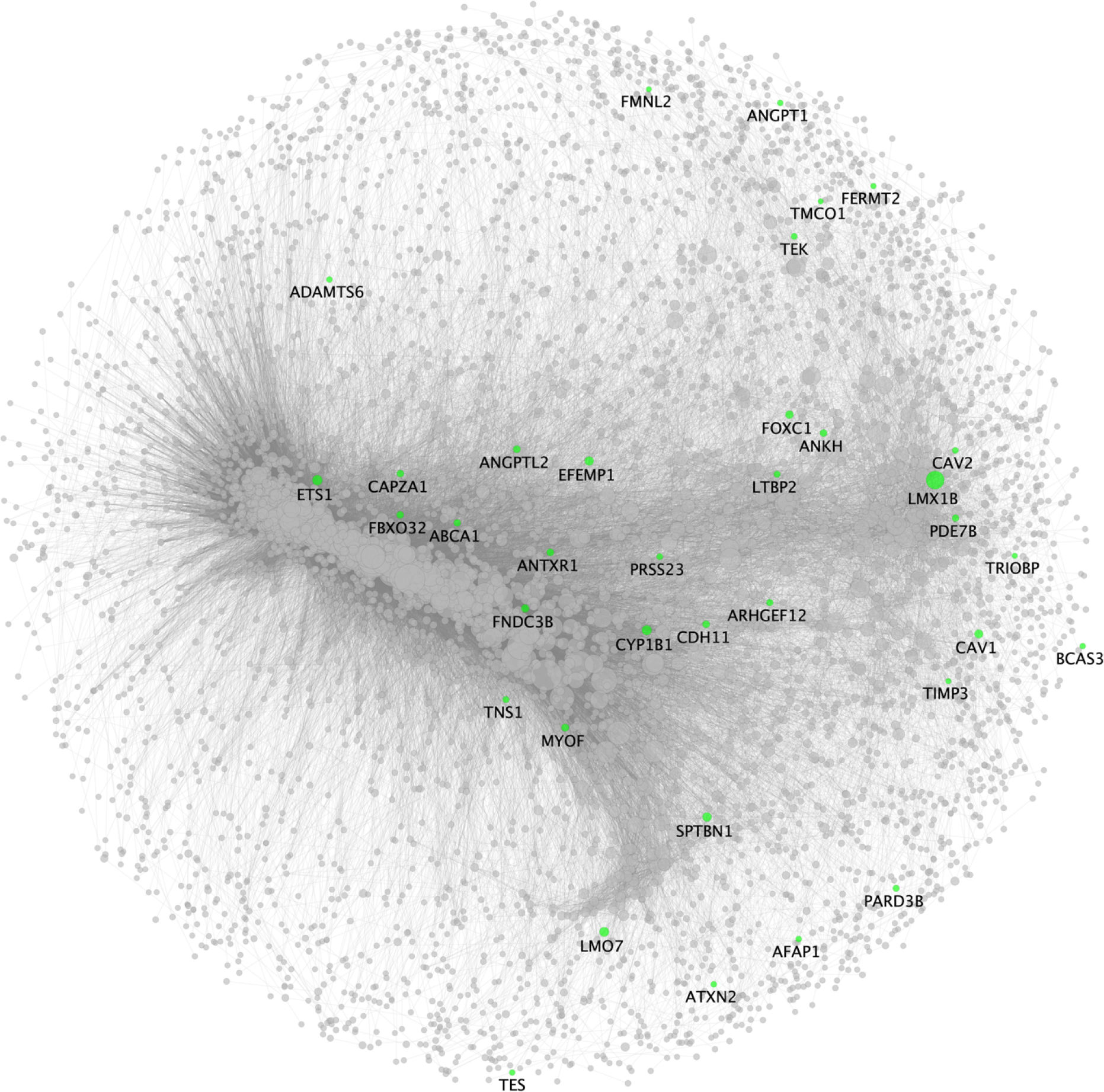
Gene expression network in non-targeting control cells. A gene expression network for non-targeting control trabecular meshwork cells was generated to highlight target genes that are normally expressed in TMCs. 30 (58%) of the 62 target genes were identified as illustrated above. Closer proximity between the genes indicate similar degrees of co-expression and the size of the node corresponds to the node centrality of PAGErank.^75^

*TAC1* has previously been shown to be upregulated in specific *MYOC* mutations.^33^ *LCE1C* (Late Cornified Envelope 1C), however, has been predicted to be involved in keratinisation and has no links to glaucoma or trabecular meshwork function in the literature. The *MYOC* knockout also induced upregulation of *SPP1,* which was not significantly present in *CAV1* or *ABO*. *SPP1* (Secreted phosphoprotein 1) encodes for a protein primarily involved in osteoclast function but has been further shown to act as a cytokine by enhancing the production of interferon-gamma and interleukin-12. Previous transcriptome analysis has shown *SPP1* is highly expressed during trabecular meshwork differentiation.^34^ Furthermore, *SPP1* is involved in retinal ganglion cell survival in-vitro via secretion by Müller cells.^35^.

Cluster eight contained four genes with a similar DEG profile; *ANGPT2, PKHD1, TNS1,* and *TXNRD2. ANGPT2* encodes for angiopoietin and is critically involved in the development of Schlemm’s canal, and abnormalities in the function of this protein are linked to raised IOP.^36–38^ *PKHD1* is a gene responsible for polycystic kidney and liver disease and has been highlighted to be potentially related to POAG pathogenesis in a family-based genetic study.^39^ *TNS1* encodes for tensin 1, which supports plasma membrane adhesion to the extracellular matrix. However, there has been no previous association with glaucoma or TM function. *TXNRD2* encodes for a thioredoxin reductase 2, which is involved in redox homeostasis and has also been associated with developing POAG.^12^ There were five significantly upregulated DEGs (*CXCL11, CST1, LCE1C, OASL, CD70*) and three downregulated DEGs (*STEAP4, CCN5, C1orf87)*. However, none of these genes has previously been attributed to the pathogenesis of POAG.

Cluster fourteen contained six genes, all with similar DEG profiles; *CAPZA1, KALRN, LMO7, PLEKHA7, GNB1L,* and *TEX41*. *CAPZA1* is involved in cytoskeletal structure via interactions with F-actin with no association with POAG. *KALRN* encodes for a protein crucial to the development of Huntington’s disease, however, has not been previously linked to POAG. *LMO7* may be involved in protein-protein interactions and has been hypothesised to contain a risk locus for developing POAG.^40^ *PLEKHA7* encodes for pleckstrin homology domain containing A7, which is primarily responsible for cell adherence and whilst being highly associated with angle-closure glaucoma^41^, has no previous association with POAG. *GNB1L* encodes for G-Protein Subunit Beta-1 Like and is involved in the formation of protein complexes. Variations in *GNB1L* have been demonstrated across geographically-distinct populations and have been associated with IOP variation, and have been speculated to be involved in the development of POAG^42^. Finally, *TEX41* is an RNA gene affiliated with the long-non-coding RNA class and also has no previous correlation with POAG. There were two key DEGs identified in this cluster, *MT1G* (upregulated) and *STEAP4* (downregulated); however, there was no previous association with POAG among these DEGs.

### Key morphological features

A heatmap was constructed showing gene knockouts with a difference of >1.5 or <-1.5 compared to non-targeting controls (p-value <1e-40) **(Figure 7)**. This identified some key genes of interest that had particular morphological changes. The *TEK* knockout group particularly showed an increase in the outputs referring to nuclear granularity identified with mitochondrial stain **(Figure 8A)**. The *LTBP2* knockout was the only group to show a significant increase in the mean nuclear intensity as well as the standard deviation nuclear intensity **(Figure 8B)**. Finally, *TRIOBP* and *TMCO1* both showed similar increases in actin / cell membrane granularity and intensity which was greater than any other groups; *TRIOBP* and *PLEKHA7* also demonstrated similar morphological profiles with similar degrees of feature increase across mitochondrial and actin / cell membrane channels **(Figure 8C)**.

**Figure 7:**
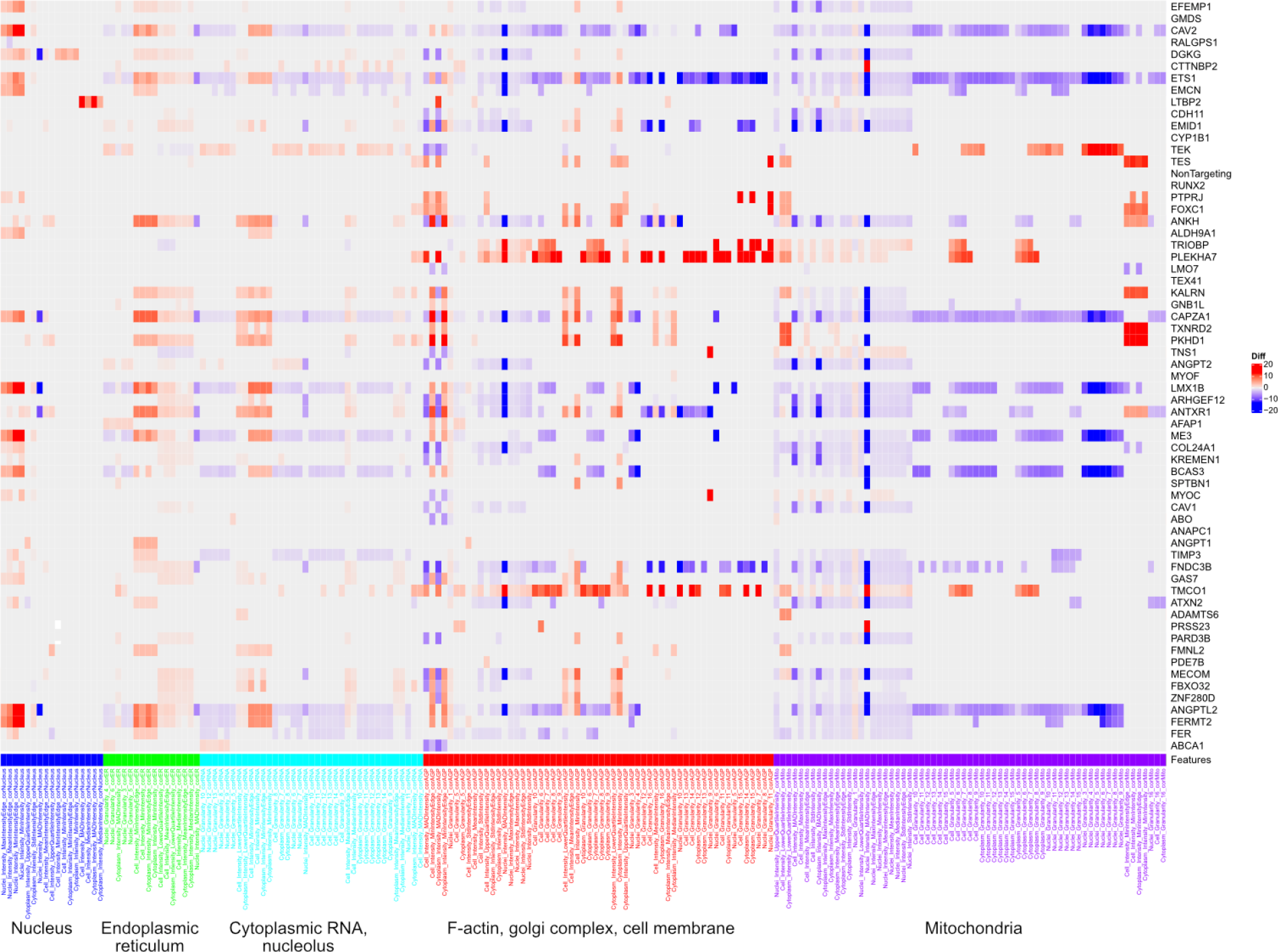
Heatmap displaying variation in TMC morphological features for knockout of genes at IOP-associated loci. The X-axis shows morphological features extracted by CellProfiler grouped by organelles of the same fluorescent channel and the Y-axis lists all gene knockouts. Red on the heatmap refers to increase in a particular morphological feature (eg. nuclear intensity) whereas blue refers to a decrease. Features extracted are based on pixel intensities and calculations based on area and appearance.

**Figure 8:**
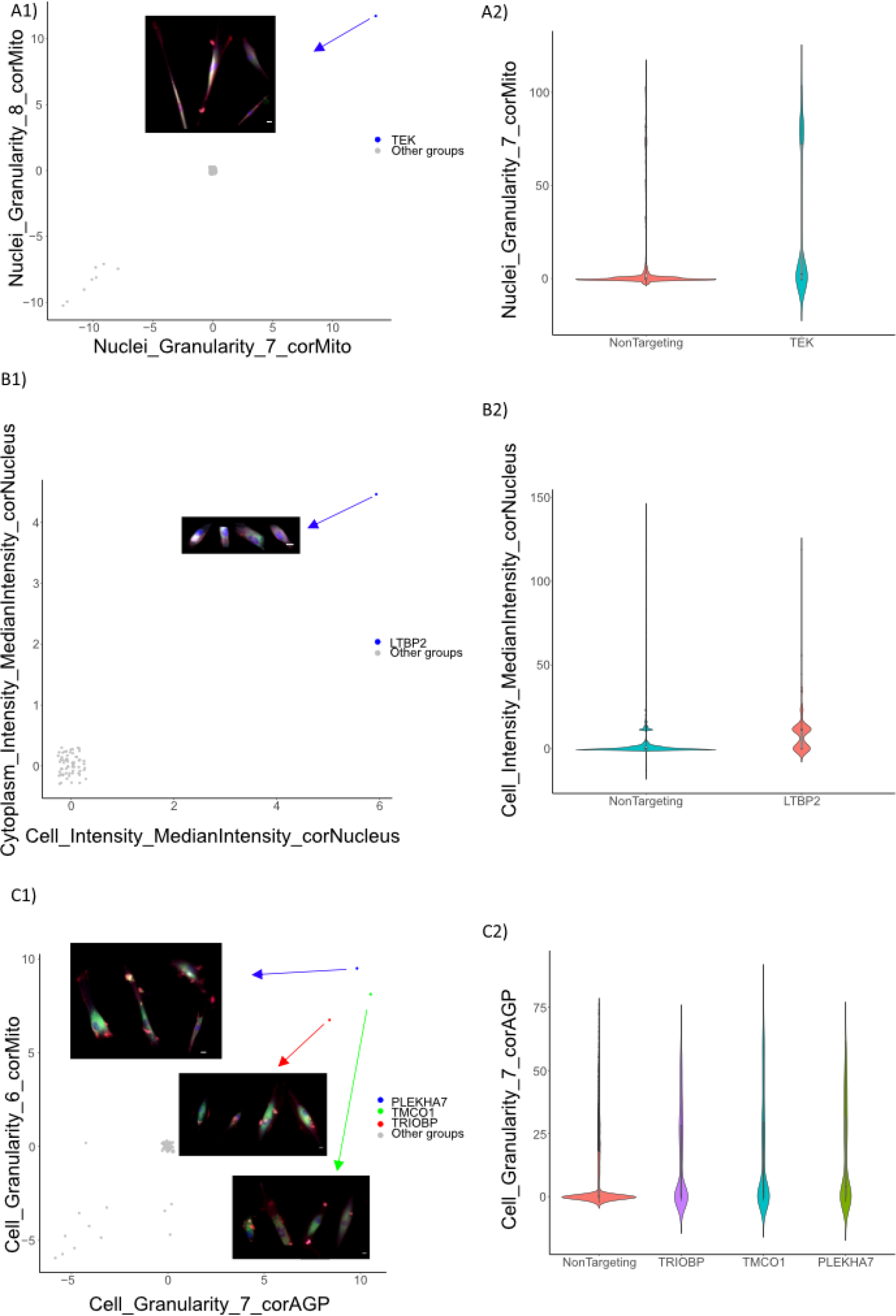
Morphological features of key gene groups of interest: A: *TEK*; B: *LTBP2*; C: *PLEKHA7*, *TMCO1*, and *TRIOBP*. Panels display the 1: Scatter plots of the mean value of each group of features (Jitter was applied to avoid overplotting). Lane2: The distribution of value of features of each group. Scale bar: 5μm

## DISCUSSION

GWAS have uncovered a large range of novel loci associated with many complex traits.^43–46^ With such significant amounts of data generated from these studies, the challenge is posed as to how to efficiently identify the most relevant gene(s) at each implicated locus.^47^ A thorough understanding of the disease is required to identify new pathological pathways and thus; new therapeutic interventions. This study sought to investigate the effects of IOP-associated gene knockouts on the morphological and transcriptome profiles of primary human TMCs. Applied in the context of POAG, this study aimed to identify genes associated with IOP to highlight potential TMC dysfunction with the goal of distinguishing new therapeutic pathways for drug discovery.

In the gene knockout groups with genes related to congenital glaucoma, *GMDS, FOXC1, MYOC, LTBP2, TEK,* and *CYP1B1,* no distinct patterns were observed in the transcriptome of these gene knockout groups. Similarly, the morphological profiles of these gene knockout groups demonstrated minimal change from non-targeting control groups. This highlights that these genes may be more involved in the development of the trabecular meshwork rather than the maintenance and as such, knocking out these genes may not significantly affect gene function or cellular morphology. The reason these gene knockouts may not illustrate a significant transcriptomic or morphological variation is that many of these genes are primarily involved in the development of the trabecular meshwork and may have a lesser role in the maintenance. Hence, a reduced response may be seen in adult human trabecular meshwork cells. Furthermore, some of these gene mutations may be gain-of-function and therefore will not exhibit a pathological response in gene knockout experiments. For example, it has been previously highlighted that genes such as *FOXC1* are primarily involved in the development of the trabecular meshwork and ocular anterior segment with mutations associated with congenital glaucoma and anterior segment dysgenesis.^48^ Similarly, studies have shown that *CYP1B1* and *LTBP2* are also involved in modulating ocular development with mutations resulting in abnormal development of the trabecular meshwork and anterior ocular circulation.^49, 50^ *MYOC* mutations are typically gain-of-function resulting in misfolded proteins inducing endoplasmic reticulum stress and extracellular matrix dysfunction.^51^ As such, loss of function *MYOC* mutations have been shown to not cause disease.^52^

When looking at gene groups of multi-gene loci, we expected to see a distinct pattern of gene expression and morphology in individual genes when compared to the control group in order to prioritise a pathogenic gene. Four of the seven knockouts were able to be prioritised with *ANGPTL2, LMX1B, CAV1,* and *EMID1* showed a higher degree of transcriptomic and morphological variation from non-targeting control cell lines than other genes associated with a given multi-gene locus. The remaining multi-gene groups all clustered separately to non-targeting controls and showed similar levels of transcriptomic and morphological variation; thus making gene prioritisation difficult. These findings allow for resolution of multi-gene loci which may contribute to the regulation of IOP and pathogenesis of POAG. Furthermore, this study presents a novel approach to resolving loci identified via GWAS with multiple potential genes candidates; a known challenge of GWAS interpretation.^44^

Hierarchical clustering was utilised to identify potential genetic networks of similar genes contributing to IOP physiology. Three clusters containing between three and five distinct gene knockouts produced similar DEG patterns indicating a potential interaction between these genes and thus; a genetic network contributing to IOP physiology and the pathogenesis of POAG. When analysing individual DEG expression across gene knockouts, it was noted that genes related to matrix metalloproteinases and interferon-related proteins were significantly up-or down-regulated. Matrix metalloproteinases have a distinct footprint of evidence showing a role in the pathogenesis of POAG.^25–27, 53, 54^ However, interferon-alpha and interferon-induced proteins have minimal previous associations with POAG potentially highlighting this as a novel pathological pathway in disease progression. Of note, *IFIH1* was the only interferon-related gene identified in DEG analysis which has been associated with glaucoma in literature. Mutations in *IFIH1* have been associated with Aicardi-Goutières syndrome and Singleton-Merten syndrome, both of which have similar overlapping features and are associated with glaucoma.^55–60^

One of the limitations of this study is that only trabecular meshwork cells have been investigated *ex vivo*, however there are several other ocular structures implicated in POAG such as the ciliary body, and Schlemm’s canal.^4^ This highlights that further investigation could be carried out to investigate the roles of IOP-related genes in cells of other areas of the eye. A further limitation of this study is that the CRISPR gene knockout results in unpredictable effects on gene function ranging from downregulation to complete silencing.^61, 62^ This may not be an accurate representation of the genetic complexity of POAG as many genes may have abnormally functioning to varying degrees from gain-of-function to deletions with complete silencing. One such example is *MYOC* in which gain-of-function mutations result in accumulation of misfolded myocilin and obstruction of the trabecular meshwork, leading to impaired outflow of aqueous humour and raised IOP.^30, 52, 63, 64^ On the contrary, other studies have highlighted a link between particular genomic deletions and POAG.^65^ Furthermore, minimal association between a gene mutation and a particular disease does not rule the gene out of playing a role in developing novel therapies. In the case of neovascular age-related macular degeneration, the genetic contribution of vascular endothelial growth factor is minimal, yet anti-VEGF agents have been remarkably successful at controlling disease.^66–68^ This highlights that phenotypic effects from strong inhibitory interventions (eg. CRISPR knockout) may be observed despite minor genetic contributions made by a particular gene. Finally, in our study we knocked out a single gene to investigate its effect on the pathogenesis of POAG and IOP regulation. However, disease processes are often contributed to by a network of genes all functioning in unison indicating that the knockout of a single gene may be insufficient to reproduce the complete disease phenotype.^69^

In the field of modern genetics, this study highlights a high-throughput approach to investigating the roles of genetic variants in disease pathogenesis. GWAS and other genomic association methods have become increasingly accessible and powerful due to cost reductions and improved computational capacity. The investigation of genetics requires quantitative analysis from multiple avenues (such as transcriptomics and morphological profiling) to fully investigate the complexities of cell biology in disease processes. This allows for the identification of genetic components of disease and thus new potential therapeutic avenues.

### CONCLUSION

In summary, this work is the first time that high-throughput multiplex morphological profiling (CellProfiler) has been combined with scRNA-seq analysis. Together, these platforms have uncovered unifying pathways involved in the homeostasis of TMCs, variation in IOP, and the pathogenesis of glaucoma. Robust pipelines have been generated to create transcriptomic and cell morphology profiles. These results demonstrate that gene perturbation can be reflected in the cell morphology with corresponding regulatory pathways, and as a consequence, this resource further improves our understanding of gene function in disease. This comprehensive transcriptomic and morphological dataset of trabecular meshwork cells represent the largest functional follow-up of genes implicated through GWAS to date. In the gene expression comparison, different cell types may be grouped according to their transcriptome patterns^70^, and the influence of the non-normal distributions and outliers may be minimized. For the cell morphology, using the median value of each feature, and adding features’ dispersion and covariances to the profiles may increase the hit rates and reliability in finding positive genes related to the disease.

## METHODS

### Cell culture

Primary human TMCs were isolated from donors through the Lions Eye Bank at the Royal Victorian Eye and Ear Hospital (ethics approval reference: 13-1151H) before being cryopreserved and delivered frozen to the Menzies Research Institute. TMCs were thawed and cultured in Dulbecco’s Modified Eagle Medium (DMEM, Gibco, 11965118) with 10% foetal bovine serum (FBS, Gibco, 16000044), and supplemented with 0.5% antibiotic-antimycotic (Gibco, 15240-062). The culture medium was changed as per local protocol after 72 hours or when cells reached 80% confluence. All cell lines were cultured at 37℃ with 5% CO_2_ in the incubator. Each fortnight cell lines were tested for mycoplasma using the PCR Mycoplasma Test Kit (PromoKine, PK-CA91-1096).

### Cloning and validation of the single-vector CROPseq system

To generate a single-vector system CROPseq plasmid expressing both SpCas9 and sgRNA(CROPseq-EFS-SpCas9-P2A-EGFP; Addgene #99248), the EF1a promoter in the CROPseq-Guide-Puro 124 (Addgene plasmid # 86708) was replaced with the EFS promoter to drive the expression of SpCas9 using the Gibson Assembly method (NEBuilder HiFi DNA Assembly master mix). The EFS-SpCas9-P2A fragment was amplified from lentiCRISPRv2 125 (Addgene plasmid # 52961) using Q5 high-fidelity DNA polymerase. The puromycin resistance gene was then subsequently replaced with EGFP using an amplified fragment from the pMLS-SV40-EGFP plasmid 126 (Addgene plasmid # 46919). The expression and activity of the single-vector CROPseq plasmid was tested by cloning in a sgRNA targeting the DNMT3B (sgRNA sequence: CAGGATTGGGGGCGAGTCGG) or LacZ control gene (sgRNA sequence: TGCGAATACGCCCACGCGAT) using Gibson Assembly method and transformed into NEBStable bacteria (NEB) as outlined by Datlinger and colleagues^19^ and tested in HEK293A cells (Life Technologies). EGFP expression was visualised using the Eclipse Ti-E inverted fluorescence microscope (Nikon). The cleavage activity of the SpCas9 was measured through the indel formation using SURVEYOR assay (Integrated DNA Technologies). Briefly, genomic DNA was extracted (QIAamp DNA mini kit; Qiagen) from HEK293A cells transfected with CROPseq-EFS-SpCas9-P2A-EGFP DNMT3B sgRNA plasmid using Fugene HD (Promega). PCR fragment for SURVEYOR assay was amplified using Q5 high-fidelity polymerase using the primers F: 5’-CAAGAGCATCACCCTAAGAATGC-3’ and R: 5’-GTTGTCAGAGACTCTCCCCAAAG-3’ from Datlinger et al.^19^. Q5 PCR conditions were as per the manufacturer’s protocol with the following thermocycling conditions: 98°C 30 secs; 35 cycles of 98°C 10 secs, 71°C 30 secs, 72°C 15 secs; 72°C 2 mins. PCR products were gel purified using the QIAquick gel extraction kit (Qiagen). 200ng of purified PCR product was used in the SURVEYOR assay as outlined in the manufacturer’s protocol.

### Confirmation of sgRNA sequence via Sanger sequencing

In total, 134 sgRNAs sequences were designed to generate the 67 trabecular meshwork cell lines (124 sgRNAs for 62 genes and 10 sgRNAs for human non-targeting control, 2 sgRNAs for each cell line) (**Supplementary Table 1**). Each of the sgRNAs was cloned into CROPseq-Guide-pEFS-SpCas9-p2a-puro backbone (Addgene: #99248). The sequences of all sgRNAs templates were confirmed by in-house Sanger sequencing. Firstly, each template was amplified by the BigDye Terminator Cycle v3.1 Sequencing kit (Applied Biosystems, 4337454). The 10 μl reaction system contained 1 μl template, 1 μl 10μM primer, 0.25 μl Reaction Mix, 1.75 μl 5X Sequencing Buffer, and 6 μl nuclease-free water. Cycling was performed using the following program: initial polymerase activation for 1 minute at 96°C and 25 cycles of amplification (denaturation for 10 seconds at 96°C, annealing for 5 seconds at 50°C, and extension for 4 minutes at 60°C), then held at 15°C. Samples were purified with the CleanSEQ kit (Beckman Coulter, A29151) following the Agencourt CleanSEQ Dye-Terminator Removal protocol. Briefly, 10 μl of vortexed CleanSEQ reagent and 42 μl of 85% ethanol was added to each 10 μl sample and gently mixed. The sample was placed on the 96-well magnetic plate for 3-5 minutes until the magnetic beads formed a ring and the solution was clear. The supernatant was removed and samples were washed twice with 100 μl 85% ethanol with 30 seconds of incubation and then air dried for five minutes. Lastly, 30 μl nuclease-free water was added to each sample and incubated for 3-5 minutes on the magnetic tray to elute the purified DNA. Next, 15 μl of purified cycle sequencing product was added to the sequencing plate, then denatured by incubating at 95°C for 5 minutes.

Sequencing Genetic Analyzer 3500, Applied Biosystems) was undertaken using the default program for 850bp DNA length. Finally, the online alignment tool MAFFT (version 7) was used to confirm whether the sequences of all the 134 sgRNAs were matched with reference sequences.

### Single-Cell RNA Sequencing

The cells were recovered in culture medium and single-cell capture was performed at the Garvan-Weizmann Centre for Cellular Genomics. Single-cell suspensions from different wells were pooled, centrifuged and resuspended in DPBS containing 1% BSA (Sigma-Aldrich, A8806-5G), and filtered by 37 μm strainer (STEMCELL, 27215). The estimated number of cells in each well in the Chromium chip was optimized to capture about 16,000 cells. The Chromium library was then generated following the protocol of the Chromium Single Cell 3’ v2 Library (10X Genomics). Briefly, individual cells were allocated into nanoliter-scale Gel Bead-in-EMulsions, in which the bead carries the primers containing a read 1 primer sequence, a 16 nt 10x barcode, a 10 nt Unique Molecular Identifier, and a poly-dT primer sequence. A barcoded, full-length cDNA was produced from each poly-adenylated mRNA after incubation with the Gel Bead-in-EMulsions. llcDNAs were pooled and amplified by PCR. In the library construction, P5, P7, a sample index, and read 2 primer sequences were added to each of the cDNA by End Repair, A-tailing, Adaptor Ligation, and PCR. The region of P5 and P7 allowed the library fragment to attach to the flow cell surface during the sequencing.

Read 1 and read 2 sequences are standard Illumina sequencing primer sites used in paired-end sequencing. Then part of the library samples were sequenced on an Illumina NovaSeq 6000 system using the S4 flowcell with a read depth of 16,785 reads per cell resulting in a mean number of RNA features of 4,195 per cell. Following this, the cell UMI-sgRNA sequence in the NGS library was also amplified and sequenced on an Illumina MiSeq-based sequencing.

### Cell Painting Immunohistochemistry

For each group, 4.0×10^3^ puromycin-selected TMCs were seeded to 96-well plates by fluorescence-activated cell sorting (FACS) via a Beckman Coulter MoFlo Astrios EQ with three replicates of each knockout group allocated at random. The whole experiment was performed in three batches of TMCs, thus, nine wells of cells were captured for each gene knockout group. The plate layout can be assessed on GitHub. TMCs were stained and fixed 48 hours after FACS following the CellPainting protocol.^20, 21^ TMCs were washed three times with HBSS without final aspiration and then sealed with parafilm. All 96-well plates were kept at 4°C in the dark before imaging.

### Automated image acquisition

Images were captured at 20X magnification in Phase Gradient Contrast (PGC), and five fluorescent channels, DAPI (385/465 nm), AF488(470/517 nm), AF514 (511/543 nm), AF594 (590/618 nm), AF647 (625/668 nm) on ZEISS Celldiscoverer 7 system. In each well, 25 sites were imaged, with autofocus in the DAPI channel as the reference.

### Morphological image feature extraction

CellProfiler (Version 3.1.9) was used to locate and segment the cells for single-cell feature extraction. The pipelines in CellProfiler were set up to correct uneven illumination, flag aberrant images and identify the nuclei from DAPI channel and the entire cell from AF594 channel, then measure the features of the size, shape, texture, intensity, and the local density of the nuclei, cell and cytoplasm.

### Establishing the CellProfiler pipeline

The CellProfiler image processing pipeline consists of three parts; illumination correction, quality control and image analysis. The illumination correction pipeline begins by improving fluorescence intensity measurement followed by the quality control pipeline to identify and exclude aberrant images such as unfocussed images and debris. To identify cell components, the nucleus was defined as the primary object with the cell body defined as the secondary object, and the cytoplasm as the tertiary object. Subsequently, the features of size, shape, granularity, colocalization, local density, and textures were measured, and the data was saved in an SQLite database. Image analysis was carried out on a Nectar (The National eResearch Collaboration Tools and Resources project) Cloud workstation instance.

### Data Curation and Analysis

Data preparation was performed using R (Version 3.6.3) as described by Caicedo et al.^71^, which included feature transformation, normalization and batch-effect correction. Firstly, all the negative controls were selected to explore the distribution of the features and the batch effects. Two transformation methods were applied, generalized logarithmic function^72^ and Box-Cox transformation.^73^ To avoid nonpositive values, generalized logarithmic function used a shrinkage strategy while Box-Cox transformation used a shift strategy.^71^ The Anderson-Darling test was performed to evaluate the normality of each feature.^74^ Next, the value of each feature was normalized by subtracting the median value of each feature from the control group and dividing by the corresponding median absolute deviation (MAD) *1.4826 in each plate, respectively. The single-cell data was aggregated by the median value of each well to create profiles of each replicate. The Spearman’s correlation was calculated for all replicates within a plate and across different plates. The replicates are selected with Spearman’s correlation score > 0.2.

### Computational analysis of single cell sequencing data

All gene knockout groups underwent hierarchical clustering and were plotted as a cluster tree. The optimal number of clusters was determined by the silhouette method. To annotate each of the clusters, the top features and tail features were extracted. The library was mapped to the GRCH38 Homo sapiens genome, and the resulting mapped counts between all samples were depth-equalized via the cellranger aggr pipeline. Peter Tran performed the MiSeq-based sequencing, and Anne Senabouth from Garvan built up the repository for the processing and analysis of single-cell RNA-seq data. In the repository, our designed gRNAs are assigned to their respective cells. Then the scRNA-seq data was loaded into R via the Seurat package (Version 3.0), and SCTransform function was used to normalise the data. All cells targeted by sgRNAs were visualised in a uniform manifold approximation and projection (UMAP) plot and were clustered with the Louvain method. The differentially expressed genes (DEGs) of each gene knockout group were selected with log2 fold change > 2 compared to the human non-targeting controls. Then a hierarchical clustering was performed on the subset of all DEGs of all gene knockout groups. The optimal number of clusters was determined by the silhouette method. DEGs to the human non-targeting controls were selected to present each cluster.

## Data & code availability

Single-cell RNA sequencing and single-cell imaging data is available at the European Bioimage Archive (Accession Numbers: S-BSST840 & S-BSST841 respectively). GitHub: https://github.com/PeterLu0403/CROP_seq_Cellpainting and https://github.com/powellgenomicslab/CROP-seq

## Acknowledgements

D.A.M., J.E.C., S.M., J.E.P, and A.W.H. are supported by the Australian National Health and Medical Research Council (NHMRC) Fellowships. X.H. was supported by the University of Queensland Research Training Scholarship and QIMR Berghofer Medical Research Institute PhD Top Up Scholarship. We are grateful for funding from Australian Vision Research; a NHMRC Program grant (1150144), Partnership grant (1132454) and the Clifford Craig Foundation.

---

**Supplementary Table 1.**
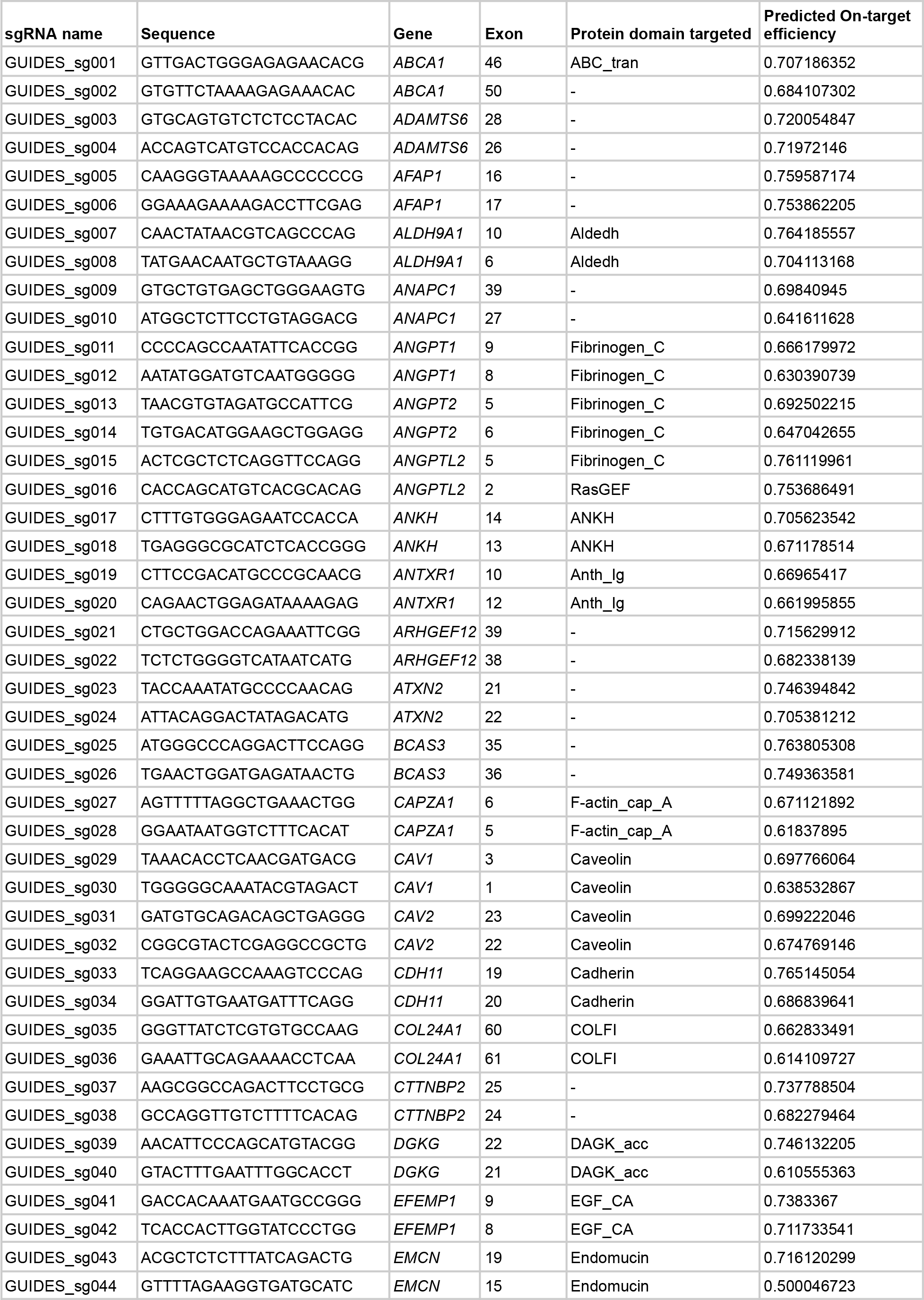
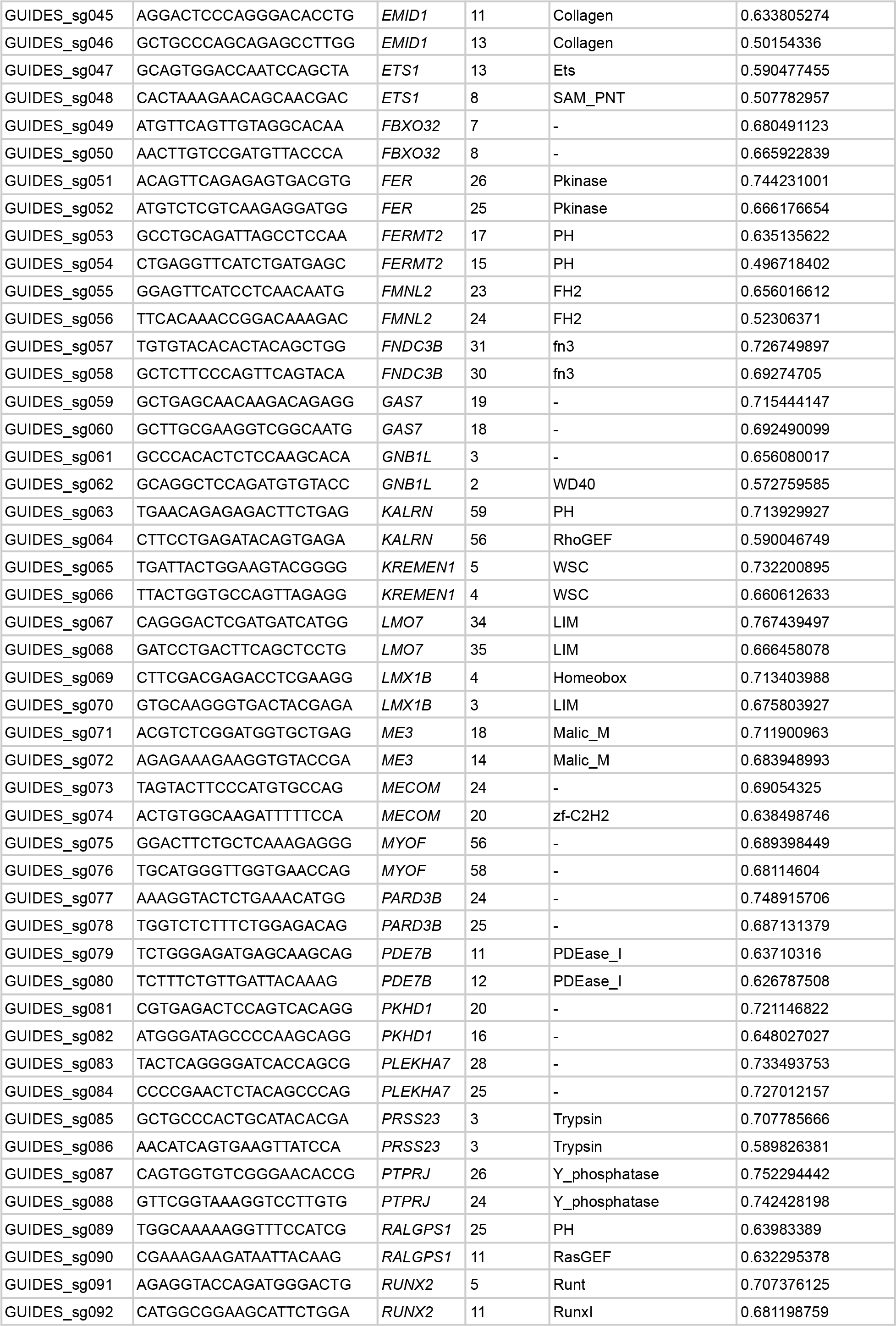
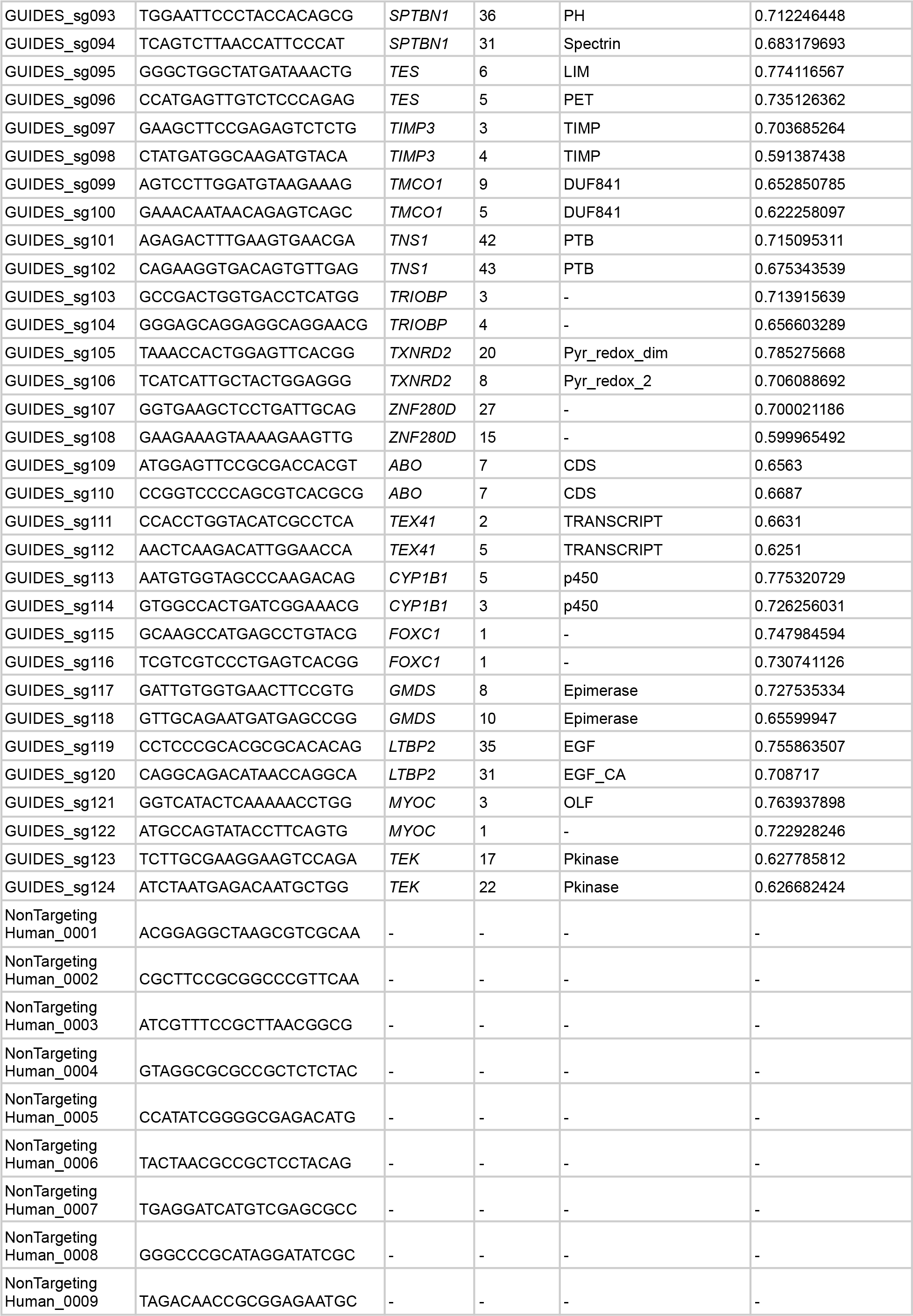
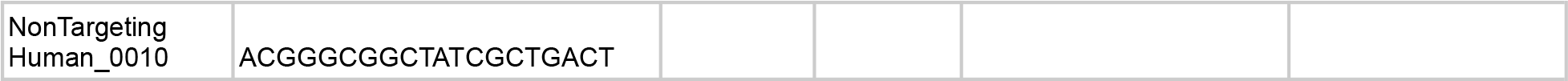

## REFERENCES

1. Kwon, Y. H., Fingert, J. H., Kuehn, M. H. & Alward, W. L. M. Primary Open-Angle Glaucoma. New England Journal of Medicine vol. 360 1113–1124 Preprint at https://doi.org/10.1056/nejmra0804630 (2009).

2. Quigley, H. A. & Broman, A. T. The number of people with glaucoma worldwide in 2010 and 2020. Br. J. Ophthalmol. 90, 262–267 (2006).

3. Boland, M. V. et al. Comparative effectiveness of treatments for open-angle glaucoma: a systematic review for the U.S. Preventive Services Task Force. Ann. Intern. Med. 158, 271–279 (2013).

4. Weinreb, R. N., Aung, T. & Medeiros, F. A. The pathophysiology and treatment of glaucoma: a review. JAMA 311, 1901–1911 (2014).

5. Wiggs, J. L. & Pasquale, L. R. Genetics of glaucoma. Hum. Mol. Genet. 26, R21–R27 (2017).

6. Fautsch, M. P. & Johnson, D. H. Aqueous humor outflow: what do we know? Where will it lead us? Invest. Ophthalmol. Vis. Sci. 47, 4181–4187 (2006).

7. Inoue, T. & Tanihara, H. Rho-associated kinase inhibitors: a novel glaucoma therapy. Prog. Retin. Eye Res. 37, 1–12 (2013).

8. Welter, D. et al. The NHGRI GWAS Catalog, a curated resource of SNP-trait associations. Nucleic Acids Res. 42, D1001–6 (2014).

9. Thorleifsson, G. et al. Common variants near CAV1 and CAV2 are associated with primary open-angle glaucoma. Nat. Genet. 42, 906–909 (2010).

10. Burdon, K. P. et al. Genome-wide association study identifies susceptibility loci for open angle glaucoma at TMCO1 and CDKN2B-AS1. Nat. Genet. 43, 574–578 (2011).

11. Gharahkhani, P. et al. Common variants near ABCA1, AFAP1 and GMDS confer risk of primary open-angle glaucoma. Nat. Genet. 46, 1120–1125 (2014).

12. Bailey, J. N. C. et al. Genome-wide association analysis identifies TXNRD2, ATXN2 and FOXC1 as susceptibility loci for primary open-angle glaucoma. Nat. Genet. 48, 189–194 (2016).

13. Kaur, K. et al. Myocilin gene implicated in primary congenital glaucoma. Clin. Genet. 67, 335–340 (2005).

14. Ali, M. et al. Null mutations in LTBP2 cause primary congenital glaucoma. Am. J. Hum. Genet. 84, 664–671 (2009).

15. Siggs, O. M. et al. Prevalence of FOXC1 Variants in Individuals With a Suspected Diagnosis of Primary Congenital Glaucoma. JAMA Ophthalmol. 137, 348–355 (2019).

16. Vasiliou, V. & Gonzalez, F. J. Role of CYP1B1 in glaucoma. Annu. Rev. Pharmacol. Toxicol. 48, 333–358 (2008).

17. Souma, T. et al. Angiopoietin receptor TEK mutations underlie primary congenital glaucoma with variable expressivity. J. Clin. Invest. 126, 2575–2587 (2016).

18. MacGregor, S. et al. Genome-wide association study of intraocular pressure uncovers new pathways to glaucoma. Nat. Genet. 50, 1067–1071 (2018).

19. Datlinger, P. et al. Pooled CRISPR screening with single-cell transcriptome readout. Nat. Methods 14, 297–301 (2017).

20. Bray, M.-A. et al. Cell Painting, a high-content image-based assay for morphological profiling using multiplexed fluorescent dyes. Nat. Protoc. 11, 1757–1774 (2016).

21. Gustafsdottir, S. M. et al. Multiplex cytological profiling assay to measure diverse cellular states. PLoS One 8, e80999 (2013).

22. Thomson, B. R. et al. A lymphatic defect causes ocular hypertension and glaucoma in mice. J. Clin. Invest. 124, 4320–4324 (2014).

23. Teixeira, L. B. C., Zhao, Y., Dubielzig, R. R., Sorenson, C. M. & Sheibani, N. Ultrastructural abnormalities of the trabecular meshwork extracellular matrix in Cyp1b1-deficient mice. Vet. Pathol. 52, 397–403 (2015).

24. Smith, R. S. et al. Haploinsufficiency of the transcription factors FOXC1 and FOXC2 results in aberrant ocular development. Hum. Mol. Genet. 9, 1021–1032 (2000).

25. He, M., Wang, W., Han, X. & Huang, W. Matrix metalloproteinase-1 rs1799750 polymorphism and glaucoma: A meta-analysis. Ophthalmic Genet. 38, 211–216 (2017).

26. Markiewicz, L. et al. Altered Expression Levels of MMP1, MMP9, MMP12, TIMP1, and IL-1β as a Risk Factor for the Elevated IOP and Optic Nerve Head Damage in the Primary Open-Angle Glaucoma Patients. Biomed Res. Int. 2015, 812503 (2015).

27. Markiewicz, L. et al. Gene polymorphisms of the MMP1, MMP9, MMP12, IL-1β and TIMP1 and the risk of primary open-angle glaucoma. Acta Ophthalmol. 91, e516–23 (2013).

28. Stone, E. M. et al. Identification of a gene that causes primary open angle glaucoma. Science 275, 668–670 (1997).

29. Goldwich, A., Scholz, M. & Tamm, E. R. Myocilin promotes substrate adhesion, spreading and formation of focal contacts in podocytes and mesangial cells. Histochem. Cell Biol. 131, 167–180 (2009).

30. Yam, G. H.-F., Gaplovska-Kysela, K., Zuber, C. & Roth, J. Aggregated myocilin induces russell bodies and causes apoptosis: implications for the pathogenesis of myocilin-caused primary open-angle glaucoma. Am. J. Pathol. 170, 100–109 (2007).

31. O’Gorman, L. et al. Comprehensive sequencing of the myocilin gene in a selected cohort of severe primary open-angle glaucoma patients. Sci. Rep. 9, 3100 (2019).

32. Wu, Z. et al. Caveolin-1 regulates human trabecular meshwork cell adhesion, endocytosis, and autophagy. J. Cell. Biochem. 120, 13382–13391 (2019).

33. Kennedy, K. D., AnithaChristy, S. A., Buie, L. K. & Borrás, T. Cystatin a, a potential common link for mutant myocilin causative glaucoma. PLoS One 7, e36301 (2012).

34. Sathiyanathan, P., Tay, C. Y. & Stanton, L. W. Transcriptome analysis for the identification of cellular markers related to trabecular meshwork differentiation. BMC Genomics 18, 383 (2017).

35. Ruzafa, N., Pereiro, X., Lepper, M. F., Hauck, S. M. & Vecino, E. A Proteomics Approach to Identify Candidate Proteins Secreted by Müller Glia that Protect Ganglion Cells in the Retina. Proteomics 18, e1700321 (2018).

36. Thomson, B. R. et al. Angiopoietin-1 is required for Schlemm’s canal development in mice and humans. J. Clin. Invest. 127, 4421–4436 (2017).

37. Thackaberry, E. A. et al. Rapid Development of Glaucoma Via ITV Nonselective ANGPT 1/2 Antibody: A Potential Role for ANGPT/TIE2 Signaling in Primate Aqueous Humor Outflow. Invest. Ophthalmol. Vis. Sci. 60, 4097–4108 (2019).

38. Kim, J. et al. Impaired angiopoietin/Tie2 signaling compromises Schlemm’s canal integrity and induces glaucoma. J. Clin. Invest. 127, 3877–3896 (2017).

39. Liu, T., Xie, L., Ye, J. & He, X. Family-based analysis identified CD2 as a susceptibility gene for primary open angle glaucoma in Chinese Han population. J. Cell. Mol. Med. 18, 600–609 (2014).

40. Gharahkhani, P. et al. Analysis combining correlated glaucoma traits identifies five new risk loci for open-angle glaucoma. Sci. Rep. 8, 3124 (2018).

41. Shuai, P. et al. Genetic associations in PLEKHA7 and COL11A1 with primary angle closure glaucoma: a meta-analysis. Clin. Experiment. Ophthalmol. 43, 523–530 (2015).

42. Shin, H.-T., Yoon, B. W. & Seo, J. H. Analysis of risk allele frequencies of single nucleotide polymorphisms related to open-angle glaucoma in different ethnic groups. BMC Med. Genomics 14, 80 (2021).

43. Uffelmann, E. et al. Genome-wide association studies. Nature Reviews Methods Primers 1, 1–21 (2021).

44. Cano-Gamez, E. & Trynka, G. From GWAS to Function: Using Functional Genomics to Identify the Mechanisms Underlying Complex Diseases. Front. Genet. 11, 424 (2020).

45. Tam, V. et al. Benefits and limitations of genome-wide association studies. Nat. Rev. Genet. 20, 467–484 (2019).

46. Visscher, P. M. et al. 10 Years of GWAS Discovery: Biology, Function, and Translation. Am. J. Hum. Genet. 101, 5–22 (2017).

47. Moore, J. H., Asselbergs, F. W. & Williams, S. M. Bioinformatics challenges for genome-wide association studies. Bioinformatics 26, 445–455 (2010).

48. Gauthier, A. C. & Wiggs, J. L. Childhood glaucoma genes and phenotypes: Focus on FOXC1 mutations causing anterior segment dysgenesis and hearing loss. Exp. Eye Res. 190, 107893 (2020).

49. Gould, D. B., Smith, R. S. & John, S. W. M. Anterior segment development relevant to glaucoma. Int. J. Dev. Biol. 48, 1015–1029 (2004).

50. Fan, B. J. & Wiggs, J. L. Glaucoma: genes, phenotypes, and new directions for therapy. J. Clin. Invest. 120, 3064–3072 (2010).

51. Donegan, R. K. et al. Structural basis for misfolding in myocilin-associated glaucoma. Hum. Mol. Genet. 24, 2111–2124 (2015).

52. Kim, B. S. et al. Targeted Disruption of the Myocilin Gene (Myoc) Suggests that Human Glaucoma-Causing Mutations Are Gain of Function. Mol. Cell. Biol. 21, 7707–7713 (2001).

53. Zhao, F., Fan, Z. & Huang, X. Role of matrix metalloproteinase-9 gene polymorphisms in glaucoma: A hospital-based study in Chinese patients. J. Clin. Lab. Anal. 34, e23105 (2020).

54. Wu, M.-Y. et al. Associations between matrix metalloproteinase gene polymorphisms and glaucoma susceptibility: a meta-analysis. BMC Ophthalmol. 17, 48 (2017).

55. Musalem, H. M., Dirar, Q. S., Al-Hazzaa, S. A. F., Al Zoba, A.-A. A. & El-Mansoury, J. Unusual Association of Aniridia with Aicardi-Goutières Syndrome-Related Congenital Glaucoma in a Tertiary Care Center. Am. J. Case Rep. 19, 500–504 (2018).

56. Bursztejn, A.-C. et al. Unusual cutaneous features associated with a heterozygous gain-of-function mutation in IFIH1: overlap between Aicardi-Goutières and Singleton-Merten syndromes. Br. J. Dermatol. 173, 1505–1513 (2015).

57. Crow, Y. J. et al. Characterization of human disease phenotypes associated with mutations in TREX1, RNASEH2A, RNASEH2B, RNASEH2C, SAMHD1, ADAR, and IFIH1. Am. J. Med. Genet. A 167A, 296–312 (2015).

58. Rice, G. I. et al. Genetic and phenotypic spectrum associated with IFIH1 gain-of-function. Hum. Mutat. 41, 837–849 (2020).

59. Xiao, W., Feng, J., Long, H., Xiao, B. & Luo, Z. H. Case Report: Aicardi-Goutières Syndrome and Singleton-Merten Syndrome Caused by a Gain-of-Function Mutation in IFIH1. Front. Genet. 12, 660953 (2021).

60. Rutsch, F. et al. A specific IFIH1 gain-of-function mutation causes Singleton-Merten syndrome. Am. J. Hum. Genet. 96, 275–282 (2015).

61. Teboul, L., Herault, Y., Wells, S., Qasim, W. & Pavlovic, G. Variability in Genome Editing Outcomes: Challenges for Research Reproducibility and Clinical Safety. Mol. Ther. 28, 1422–1431 (2020).

62. Lino, C. A., Harper, J. C., Carney, J. P. & Timlin, J. A. Delivering CRISPR: a review of the challenges and approaches. Drug Deliv. 25, 1234–1257 (2018).

63. Ueda, J., Wentz-Hunter, K. & Yue, B. Y. J. T. Distribution of myocilin and extracellular matrix components in the juxtacanalicular tissue of human eyes. Invest. Ophthalmol. Vis. Sci. 43, 1068–1076 (2002).

64. Resch, Z. T. & Fautsch, M. P. Glaucoma-associated myocilin: a better understanding but much more to learn. Exp. Eye Res. 88, 704–712 (2009).

65. Kaurani, L. et al. Gene-rich large deletions are overrepresented in POAG patients of Indian and Caucasian origins. Invest. Ophthalmol. Vis. Sci. 55, 3258–3264 (2014).

66. Solomon, S. D., Lindsley, K., Vedula, S. S., Krzystolik, M. G. & Hawkins, B. S. Anti-vascular endothelial growth factor for neovascular age-related macular degeneration. Cochrane Database Syst. Rev. 3, CD005139 (2019).

67. Kaiser, S. M., Arepalli, S. & Ehlers, J. P. Current and Future Anti-VEGF Agents for Neovascular Age-Related Macular Degeneration. J. Exp. Pharmacol. 13, 905–912 (2021).

68. DeAngelis, M. M. et al. Genetics of age-related macular degeneration (AMD). Hum. Mol. Genet. 26, R45–R50 (2017).

69. Qassim, A. et al. An Intraocular Pressure Polygenic Risk Score Stratifies Multiple Primary Open-Angle Glaucoma Parameters Including Treatment Intensity. Ophthalmology 127, 901–907 (2020).

70. van Zyl, T. et al. Cell Atlas of Aqueous Humor Outflow Pathways in Eyes of Humans and Four Model Species Provides Insights into Glaucoma Pathogenesis. Preprint at https://doi.org/10.1101/2020.02.04.933911.

71. Caicedo, J. C. et al. Data-analysis strategies for image-based cell profiling. Nat. Methods 14, 849–863 (2017).

72. Huber, W., von Heydebreck, A., Sültmann, H., Poustka, A. & Vingron, M. Variance stabilization applied to microarray data calibration and to the quantification of differential expression. Bioinformatics 18 Suppl 1, S96–104 (2002).

73. Durbin, B. P., Hardin, J. S., Hawkins, D. M. & Rocke, D. M. A variance-stabilizing transformation for gene-expression microarray data. Bioinformatics 18 Suppl 1, S105–10 (2002).

74. Marsaglia, G. & Marsaglia, J. Evaluating the Anderson-Darling Distribution. J. Stat. Softw. 9, 1–5 (2004).

75. Iacono, G. et al. bigSCale: an analytical framework for big-scale single-cell data. Genome Res. 28, 878–890 (2018).

